# Dynamic association of IκBα to chromatin is regulated by acetylation and cleavage of histone H4

**DOI:** 10.1101/2021.02.15.431217

**Authors:** Laura Marruecos, Joan Bertran, Daniel Álvarez-Villanueva, Martin Floor, María Carmen Mulero, Anna Vert, Yolanda Guillén, Sara Arce, Laura Batlle, Jordi Villà-Freixa, Gourisankar Ghosh, Anna Bigas, Lluís Espinosa

## Abstract

IκBs exert a principal function as cytoplasmic inhibitors of the NF-kB transcription factors. Additional functions for IκB homologues have been described including association to chromatin and transcriptional regulatioin. Phosphorylated and SUMOylated IκBα (pS-IκBα) binds histones H2A and H4 in the stem and progenitor compartment of skin and intestine, but the mechanisms controlling its recruitment to chromatin are largely unstudied.

We here show that serine 32-36 phosphorylation of IκBα favors its binding with nucleosomes and demonstrated that p-IκBα association to H4 is favored by acetylation at specific H4 lysine residues. N-terminal tail of H4 is lost during intestinal cell differentiation by proteolytic cleavage at residues 17-19 imposed ny trypsin or chymotrypsin, which interferes p-IκBα binding. Paradoxically, inhibition of trypsin and chymotrypsin activity in HT29 cells increased p-IκBα chromatin binding and impaired goblet cell differentiation, comparable to IκBα deletion. Together our results indicate that dynamic binding of IκBα to chromatin is a requirement for intestinal cell differentiation and provide a molecular base for the restricted nuclear distribution of p-IκBα at specific stem cell compartments.

## INTRODUCTION

Cytoplasmic IκB proteins play an essential role as negative regulators of the NF-κB pathway, which controls immune responses and inflammation from hydra to mammals (Hoffmann and Akira, 2013). Canonical activation of NF-κB is initiated by stimuli such as TNFα, IL1β or LPS and requires IκB kinase (IKK)-induced phosphorylation and subsequent ubiquitination-mediated degradation of the IκB inhibitors. Alternative nuclear functions for various IκB and IκB-like family members including IκBα (Arenzana-Seisdedos et al., 1997; Marruecos et al., 2020; Mulero et al., 2013), IκBβ (Rao et al., 2010), IκBNS (Hirotani et al., 2005) or Bcl3 (Bours et al., 1993) have been reported. The molecular mechanisms that dictate these moonlighting nuclear functions remain understudied.

Previous studies from different groups (Culver et al., 2010; Desterro et al., 1998; Hendriks et al., 2014; Marruecos et al., 2020; Mulero et al., 2013) demonstrated that a fraction of the IκBα protein is SUMOylated at the same K21 residue that, when ubiquitinated, triggers IκBα degradation. Thus, SUMOylated-IκBα is primarily protected from degradation even in its phosphorylated form and independent of stimuli such as TNFα (Mulero et al., 2013). Phosphorylated and SUMOylated IκBα (pS-IκBα) is localized in the nucleus of basal layer keratinocytes (Mulero et al., 2013) and in the intestinal crypt compartment (Marruecos et al., 2020), where stem and progenitor cells of both tissues reside, to regulate cytokine-dependent activation of a subset of polycomb repression complex (PRC) 2 target genes (Marruecos et al., 2020; Mulero et al., 2013). SUMOylated IκBα shows reduced association with NF-kB factors (Culver et al., 2010; Mulero et al., 2013), but efficiently binds the N-terminal tail of histones, in particular H2A and H4. In fact, binding of IκBα to histones precludes its subsequent association with NF-kB factors (Mulero et al., 2013).

Histones are an essential structural and regulatory element of the chromatin and they are organized in a highly stable structure called nucleosome. Nucleosomes are formed by several subunits of histone H2A, H2B, H3 and H4 tightly associated with DNA. Importantly, the N-terminal tail of histones is flexible and protrudes outward the nucleosome, which makes histones accessible to editing enzymes that decorate their tails with a high variety of post-translational modifications (PTMs). PTMs can be dynamically added and removed thus modulating gene activation, silencing, chromatin accessibility, replication and DNA repair, in part by association with a plethora of non-histone proteins including transcription factors. Prototypical examples of proteins that specifically recognize PTMs in histones are the <u>br</u>omo<u>d</u>omains (BRDs) and the BET (bromodomains and extra-terminal) family of proteins. BETs bind specific (Ac) K residues of histones in active regulatory domains such as promoters and enhancers (Filippakopoulos et al., 2012).

We here studied the molecular requirements for pS-IκBα binding to chromatin, the residues involved in pS-IκBα to histone H4 interaction, and the mechanisms that impose restricted pS-IκBα distribution in the stem cell compartments of the skin and intestine.

## RESULTS

### Hydrophobic interactions mediate p-IκBα association to histone H4

Histones are basic proteins, and their positive charges determine association with the DNA. We investigated the possibility that electrostatic forces exert a major role in the observed association of IκBα with histones. We found that full-length (FL) and two different amino-terminal IκBα fragments (aa1-200; aa20-206) robustly interacted with histone H4 even at high salt concentrations (300mM NaCl) (Figure S1A), indicating the involvement of hydrophobic forces in histones-IκBα binding. We aimed to confirm the stability of the interaction by <u>F</u>ast *P*rotein <u>Li</u>quid <u>C</u>hromatography (FPLC) analysis of IκBα and histone H4 complexes. However, histone H4 was highly degraded in this assay and was replaced by H2A (previously found to similarly bind IκBα). Chromatography analysis demonstrated that both IκBα and H2A eluted at fraction 15-17 when run together, whereas IκBα or H2A alone eluted at fractions 29-30 and 18-20, respectively. This shift strongly suggested the existence of stable complexes that were composed of equimolar amounts of IκBα and H2A proteins as determined by Coomassie Blue staining of the eluted fractions (Figure S1B). Then, we performed in vitro phosphorylation of IκBα and SUMOylated IκBα (Figure S1C) and tested the capacity of different IκBα species to bind to reconstituted nucleosome core particles (NCP) (instead of individual histones). We observed a significantly higher capacity of in vitro phosphorylated IκBα to form stable NCP-IκBα complexes compared with non-phosphorylated IκBα (Figure 1A). Of note that band retardation imposed by p-IκBα and pS-IκBα was not significantly different likely due to loss of the SUMO chain during the experimental procedure. As an additional control, NCP-IκBα complexes were prevented or displaced by the addition of anti-IκBα in the binding reaction (Figure 1B).

**Figure 1.**
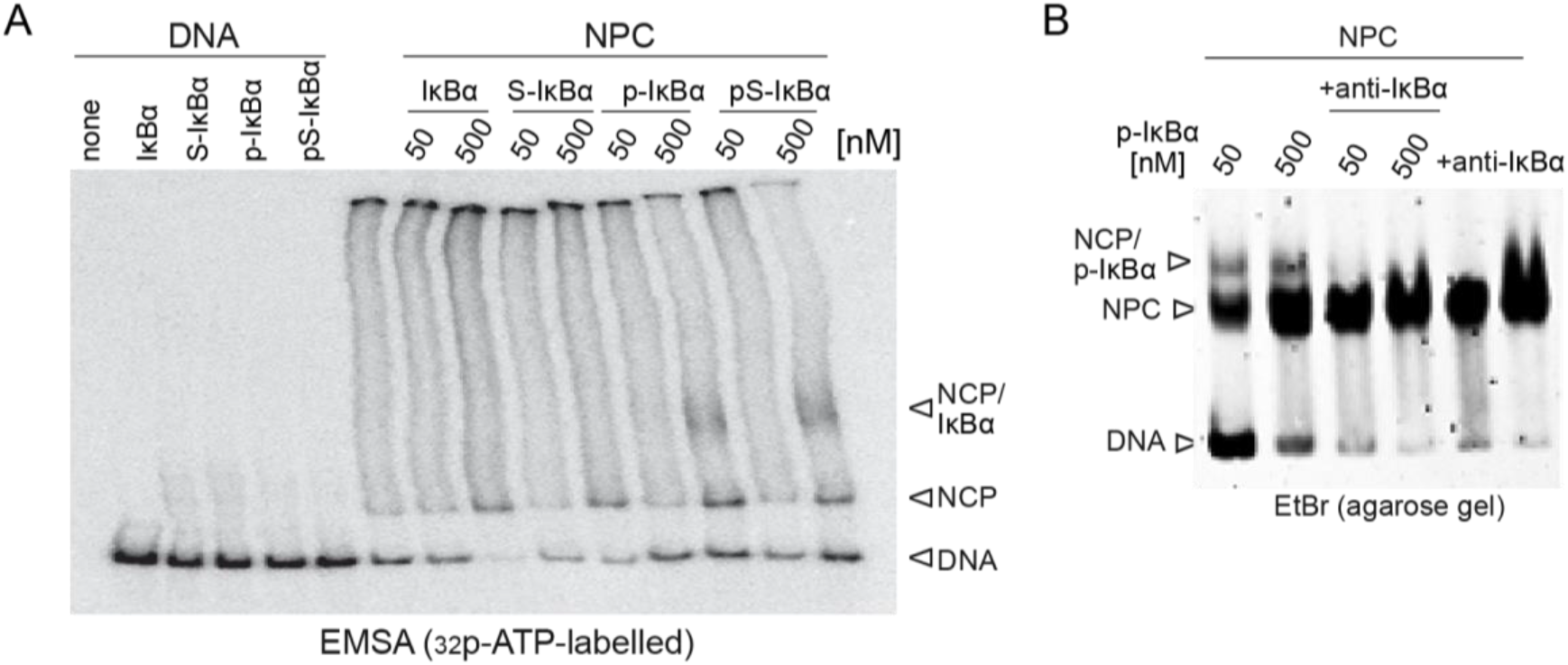
Specific interaction of nucleosome particles with phosphorylated IκBα. (A) Electrophoretic analysis (under non-denaturing conditions) of the association between the indicated IκBα species association and reconstituted nucleosome core particles (NCP). (B) Electrophoretic analysis in agarose gels of in vitro generated p-IκBα and reconstituted nucleosome in absence or presence of anti-IκBα antibodies.

Our results indicate that amino acids 1-200 of IκBα are required for H4 binding, which is independent of electrostatic forces and favored by IκBα phosphorylation.

### Preferential association of IκBα with acetylated histone H4 in vivo

We previously demonstrated that pS-IκBα binds the N-terminal tail of histones H2A and H4 and identified acetylated K12 and K16 of H4 as preferential motifs for pS-IκBα binding in vitro (Mulero et al., 2013). To further investigate pS-IκBα binding specificity, we generated biotinylated peptides of the N-terminal H4 sequence (aa1-23) including specific PTM combinations and used them to pull-down (PD) IκBα from cell extracts (Figure 2A). Both pS-IκBα (≈60 kDa) and non-SUMOylated p-IκBα (≈37 kDa) bands were consistently detected in the precipitates from non-modified H4 peptide, single K12Ac, K20Ac and double K12,16Ac peptides. In contrast, mutation of K residues to A, acetylation (Ac) of all four K residues (K5, K8, K12 and K16) or single K5Ac, K8Ac significantly reduced IκBα binding to histone H4. Comparable binding inhibition was detected in peptides containing methylated K12 and K20 (Figures 2B). Binding affinity was confirmed in PD assays using GST-SUMO-IκBα as bait and serial dilutions of histone-enriched cell extracts (see methods) with the highest signal corresponding to the H4K16Ac and H4K20Ac marks and the lowest to H4K20me2,3 (Figures 2C). These results indicate that hyperacetylation or methylation of K residues reduce IκBα to histone H4 binding, whereas H12, K16 and K20 acetylated histone binds to IκBα similar to the non-acetylated histone.

To study whether H4 acetylation favors IκBα binding to chromatin *in vivo*, we performed chromatin immunoprecipitation (ChIP) assay from HCT-116 colorectal cancer cells using the antibody against IκBα (IκBα, sc-307) and 2 antibodies against H4KAc (pan-H4KAc, Abcam ab177790 and H4K12ac, Active Motif 61527). Analysis of data demonstrated significant enrichment of IκBα peaks in acetylated chromatin regions (p<0.01) compared with random distribution of peaks from both precipitations (Figure 2D). Moreover, we detected higher accumulation of H4KAc around the TSS of IκBα-bound genes compared with non-selected gene promoters (labelled as All genes) (Figure 2E, upper panels) as well as a significant overlap between IκBα and H4KAc peaks at specific genomic loci (Figure 2E, lower panels).

**Figure 2.**
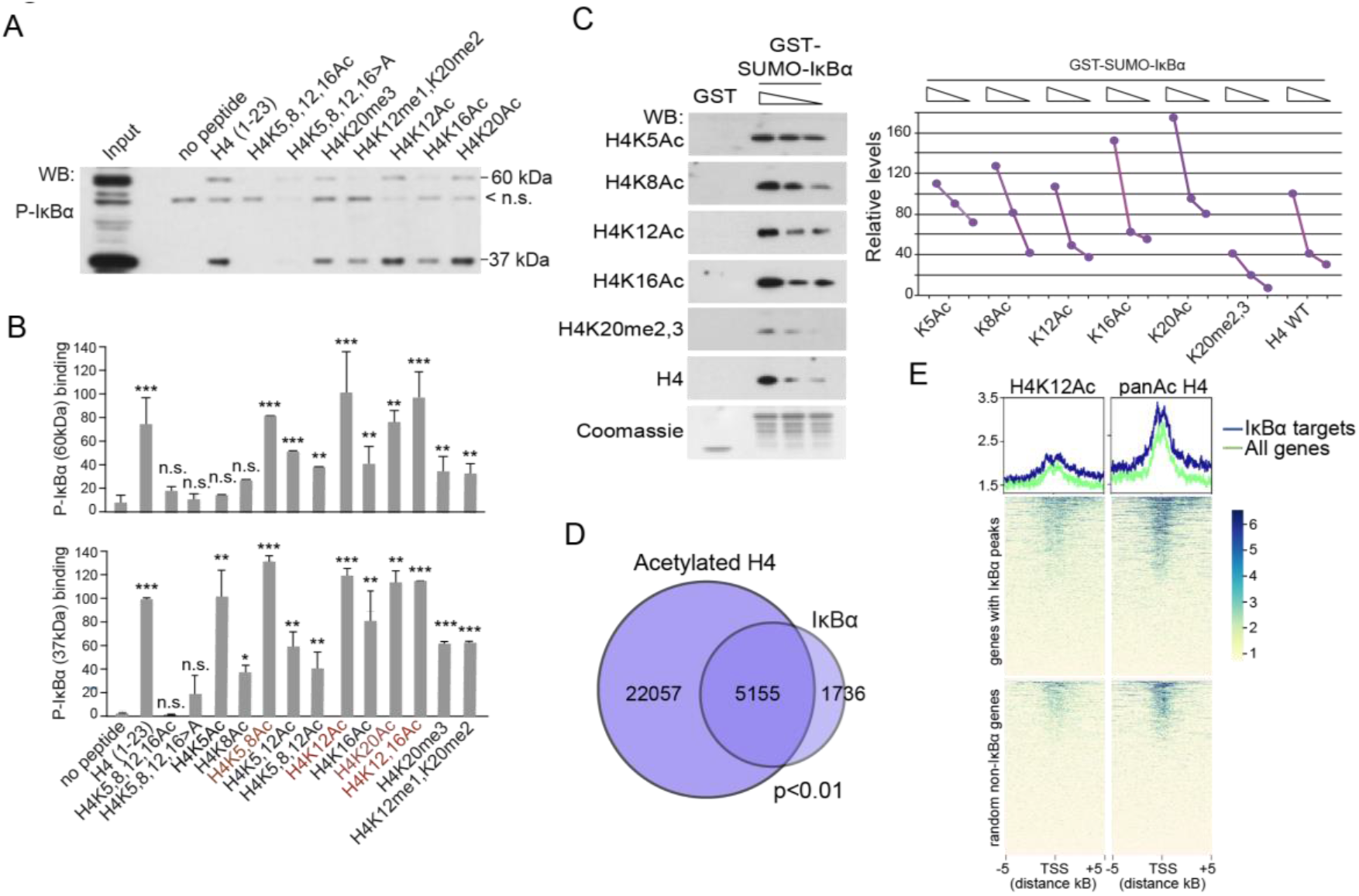
Preferential association of IκBα with N-terminal acetylated histone H4. (A) PD experiment with biotinylated peptides of H4 (1-23aa) and total lysates of HCT-116 cells. (B) Quantification of the IκBα 60 kDa band (upper panels) and the 37 kDa band (lower panels) is shown as average and s.d. of three independent experiments. (C) PD experiments with GST-SUMO IκBα fusion protein and serial dilutions of histone-enriched HEK-293T lysates. Quantification of three independent replicates is shown. (D) Venn diagrams representing the overlap between IκBα and acetylated H4 peaks obtained in ChIP-seq experiments from HCT-116 cells. (E) Peak distribution of acetylated H4 and H4K12ac obtained in IκBα target genes and non-IκBα targets relative to the transcription start site (TSS). WB in A and C are representative of three independent replicates performed. In B, *p* values were derived from unpaired two-tailed *t*-test of triplicates, ***p-value<0.001, **p-value<0.01, * p-value<0.05, n.s. no significant.

These results indicate that IκBα similarly binds acetylated histone H4 in vitro, which is precluded by K methylation, and preferentially binds acetylated H4 in vivo.

### Acetylated H4 species are restricted to stem cell compartments and lost in differentiated cells associated to histone N-tail cleavage

Previous studies from our group demonstrated that nuclear pS-IκBα is localized in the stem cell compartments of skin and intestine (Marruecos et al., 2020; Mulero et al., 2013). Our results suggest that pS-IκBα preferentially bind acetylated histone H4 in vivo (see Figures 2D and 2E). Thus, we studied the possibility that nuclear IκBα was restricted to areas containing specific H4KAc marks. Different antibodies against acetylated histone H4 labeled cells localized in the intestinal crypt compartment colocalizing with p-IκBα (Figure 3A), and including the canonical Lgr5+ ISCs (Figure 3B). Similarly, H4K12Ac was primarily detected in the keratinocytes of the basal layer of skin and the hair follicles, where progenitors and stem cells reside (Figure S2A). In contrast, H4K20me2,3, which showed the lower affinity for IκBα binding (see Figure 1A-C), was exclusively present in differentiated cells of the intestinal villi and skin (Figure 3A and S2A). The few intestinal crypt cells that contained H4K20me2,3 mark were identified as terminally differentiated Paneth cells based on their morphology and localization (Figure 3A and 3B). Compartmentalization of H4KAc and H4K20me marks was progressively reached during embryonic development (Figure S2B), which parallels the progressive restriction of nuclear IκBα distribution in the developing intestinal tissue (Marruecos et al., 2020).

**Figure 3.**
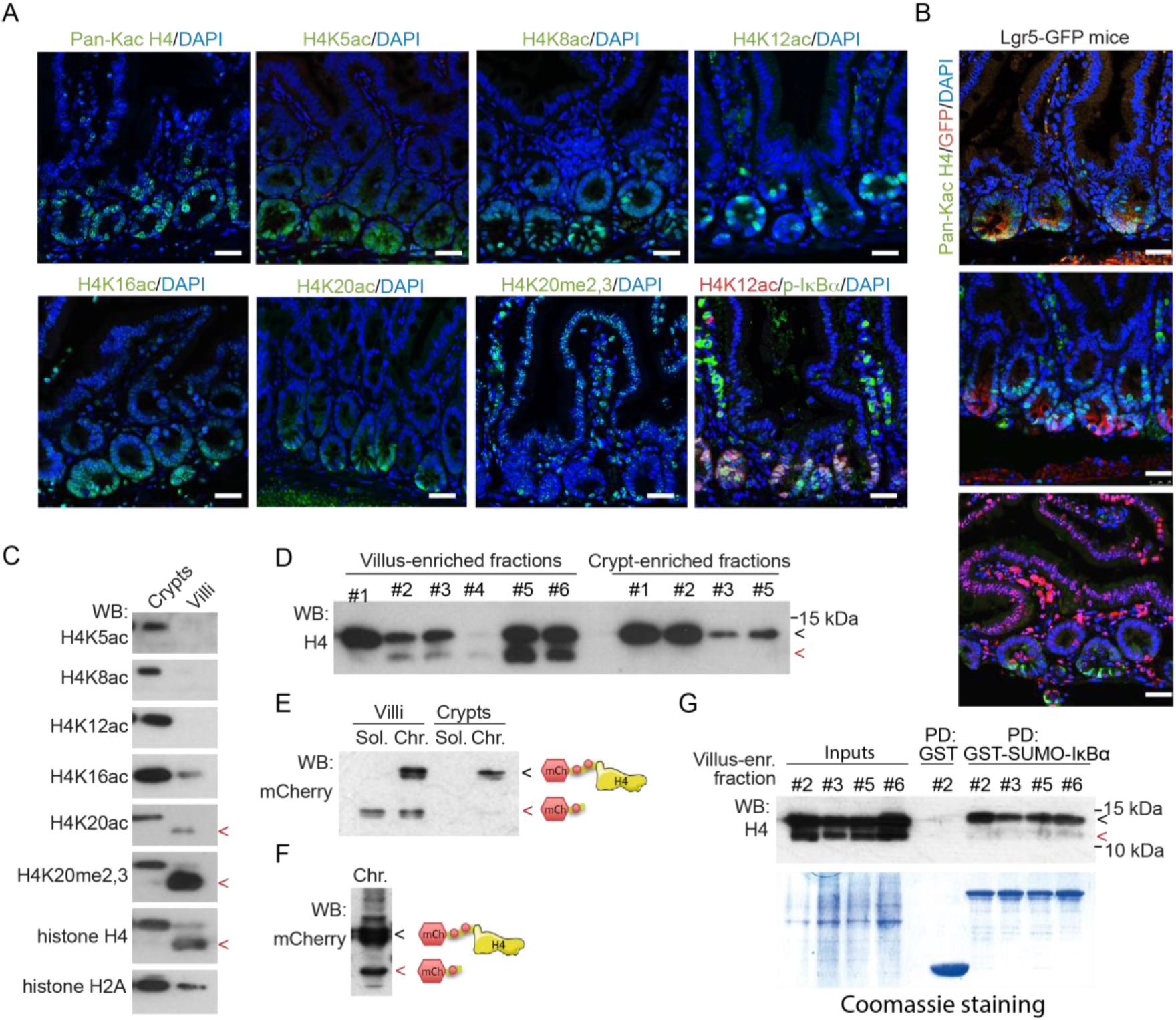
Stem cell compartments contain acetylated H4 species that are lost following histone cleavage. (A) Immunofluorescence (IF) analysis with the indicated antibodies in sections from murine small intestine of 2-month-old WT mice. (B) Double IF analysis with the indicated antibodies in sections from murine small intestine of 2-month-old Lgr5-GFP transgenic mice. (C) Western Blot analysis of chromatin extracts from isolated intestinal villus and crypt cells. Western Blot analysis of chromatin extracts from isolated intestinal villus and crypt cells in several 2-month-old WT mice (each # represents a mouse). Western Blot analysis of soluble (Sol) and chromatin (Chr) extracts from isolated intestinal villus and crypt cells in 2-month-old mCherry-H4 transgenic mice. (F) Western Blot analysis of chromatin fraction from mCherry-H4 transfected HCT-116 cells. (G) Pulldown assay using GST-SUMO IκBα protein and chromatin lysates from 2-month-old WT mice villi. Scale bars in A and B, 25μm. Red arrowheads in C-G are showing the location of truncated histone H4.

Specific distribution of acetylated histone H4 restricted to the intestinal crypts was confirmed by western blot of crypts and villus-enriched fractions (Figure 3C). Remarkably, WB analysis of total histone H4, H4K20me2,3 and H4K20KAc revealed the presence of a low molecular weight (LMW) band (red arrowheads in Figure 3C) specifically in the villus-enriched fractions. This result was confirmed in independent villus- and crypt-enriched extracts obtained from several mice (Figure 3D). The fact that the LMW band that was not recognized by the H4K5, K8, K12 and K16Ac antibodies (see Figure 3C) strongly suggested that the cleavage site of H4 mapped between K16 and K20 residues comprising the sequence KRHRK. Using a transgenic mouse line carrying mCherry fused to H4 (see methods), we confirmed that histone cleavage occurs in the intestinal villi in vivo, and involved the N-terminal tail of histone H4 leading to the release of mCherry from the H4 protein (Figure 3E). Comparable results were obtained by transfection of this construct in HCT-116 CRC cells (Figure 3F). PD experiments using chromatin extracts from intestinal villus demonstrated that IκBα specifically bound full-length histone H4 but failed to associate with truncated histone H4 lacking the N-terminal tail (Figure 3G).

Our results indicated that intestinal cell differentiation correlates with N-terminal truncation of histone H4 resulting in a cleaved histone H4 form that is unable to bind IκBα.

### Cleavage of the N-terminal tail of H4 is mediated by chymotrypsin an trypsin activity present in the intestinal villus

To address the possibility that histone H4 cleavage was due to the activity of a protease specifically contained in the differentiated intestinal cells, we incubated crypt-derived chromatin extracts (containing intact histone H4) with soluble lysates from either villus-derived or crypt-derived cells. Cleavage of histone H4 was specifically induced by incubation with villus-derived lysates (Figure 4A) further indicating that one or more proteases present and active in the differentiated cells induce histone H4 cleavage. Then, we obtained villus- and crypts-enriched cell fractions (by cell sorting based on EPHB2 levels), purified the RNA and performed RNA-seq analysis. We identified several proteases that were specifically expressed in the villus-derived RNA (Figure 4B). Bioinformatic analysis using the PeptideCutter-ExPASy (Wilkins et al., 1999) and PROSPER (Song et al., 2012) tools identified the serine proteases trypsin and chymotrypsin C as the most likely enzymes capable to cleave histone H4 in the region involving residues K16 to K20, which we have experimentally identified as the sites for cleavage. By point mutation of specific H4 residues in mCherry-H4 fusion protein, we recognized amino acids 17-19 as essential for H4 cleavage (Figure 4C), which is consistent with trypsin or chymotrypsin as responsible for this activity.

**Figure 4.**
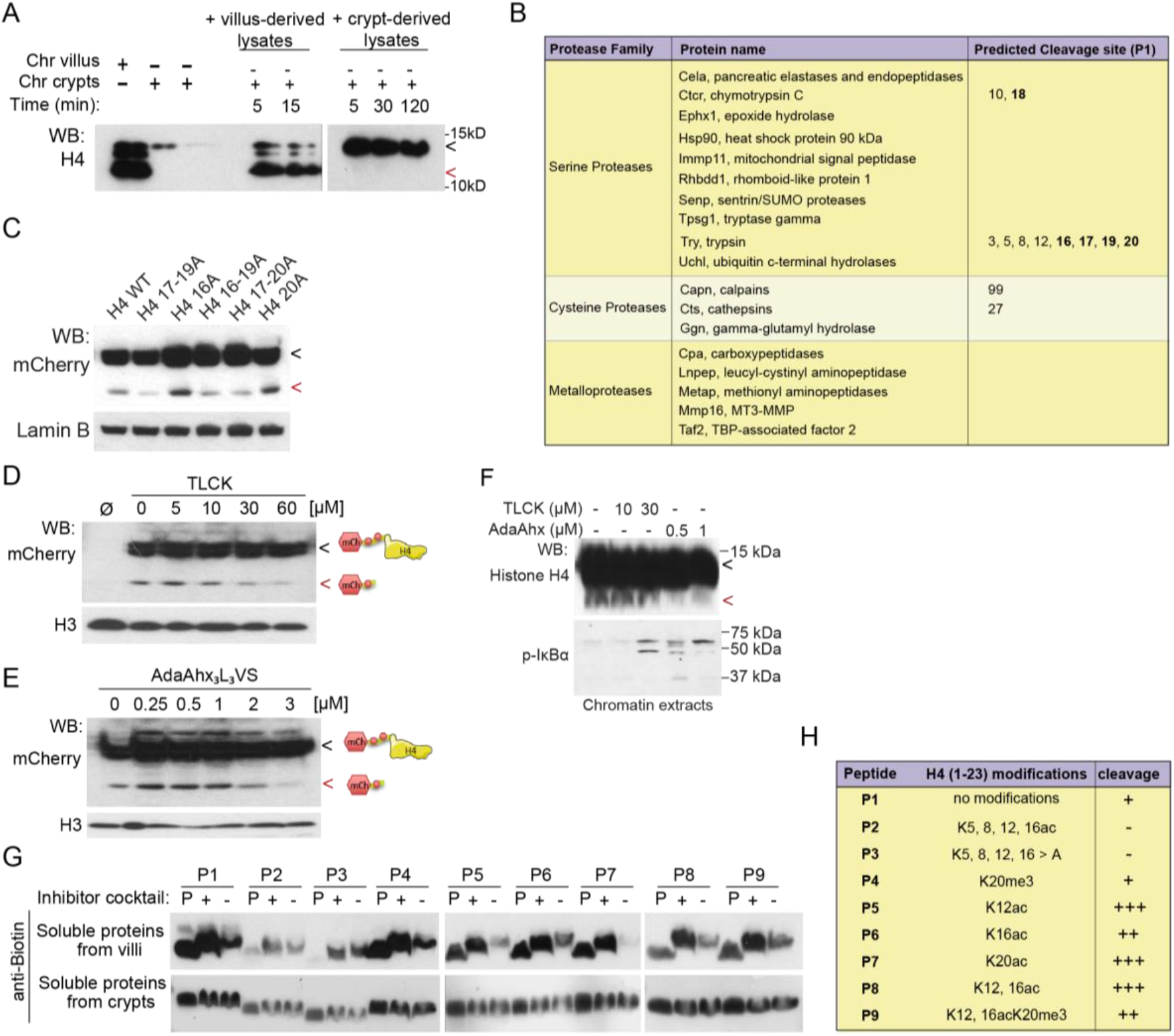
Cleavage of the N-terminal tail of H4 is mediated by chymotrypsin and trypsin activity present in the intestinal villus. (A) Western Blot analysis of the crypt-derived chromatin fractions incubated with soluble lysates from villi or crypts for the indicated times. (B) Table showing the proteases differentially expressed in the villus compartment as determined by RNA-seq analysis of intestinal crypt (purified EphB2 high cells) and villus (EphB2 negative/low) cells. Predicted cleavage sites for each protease in the histone H4 sequence was determined using the PeptideCutter-ExPASy and PROSPER software. (C) Western blot analysis of mCherry-H4 with the indicated mutations transfected in HCT-116 cells. (D, E) Western Blot analysis of mCherry-H4 transfected in HCT-116 cells treated for 16 hours with TLCK (D) or AdaAhx_₃_L_₃_VS (E) at the indicated concentrations. (F) Western blot analysis of chromatin extracts from post-confluent HT29 cells treated as indicated. (G) Western Blot analysis of a cleavage experiment incubating modified H4 peptides (P1-P9) with soluble lysates from villi or crypts in the presence (+) or absence (-) of the commercial protease inhibitors cocktail. Lanes indicated as P correspond to the control peptide without lysate incubation. Notice the electrophoretic shift of peptides incubated with villus lysates (compared with control peptides) suggestive of their binding to proteins absent from crypt lysates. (H) Quantification of the relative cleavage of specific H4 peptides incubated with villus-derived lysates from 3 independent experiments performed. Red arrowheads in A, C-F indicate truncated histone H4.

To further test whether trypsin or chymotrypsin execute histone H4 cleavage, we treated mCherry-H4 expressing HCT-116 cells with the specific inhibitors of trypsin-like proteases TLCK and AdaAhx_3_L_3_VS for 16hours. WB analysis indicated that TLCK (Figure 4D) or AdaAhx_3_L_3_VS (Figure 4E) imposed a dose-dependent inhibition of mCherry-H4 (Figures 4D and 4E) and endogenous H4 cleavage (Figure 4F). Moreover, treatment with protease inhibitors was associated with increased chromatin retention of p-IκBα (≈37 kDa) and pS-IκBα (≈60 kDa) as determined by WB of chromatin extracts (Figure 4F).

It is established that trypsin and trypsin-like proteases digest substrates containing unmodified K residues but this activity can be affected by K acetylation due to both steric effects and loss of the positive charge (Huang et al., 2015). We used different histone H4 peptides (aa1-23) to determine whether protease activity from villus-derived extracts was modified by specific K modifications. Our data indicated that villus extracts (but not with crypt-derived extracts) induced the degradation of histone H4 peptides with the only exception of the K to A mutant and the peptide with all K residues acetylated (K5,8,12,16Ac) (Figure 4G and 4H).

Together these results indicate that trypsin and chymotrypsin can induce histone H4 cleavage, and suggest that these proteases are responsible for physiologic N-terminal H4 truncation that occurs during intestinal differentiation.

### Functional impact of IκBα chromatin binding dynamics in cell differentiation

IκBα deficiency results in altered stem cell maturation and defective intestinal and skin differentiation in mice (Marruecos et al., 2020; Mulero et al., 2013). To further investigate the impact of IκBα association to chromatin in intestinal differentiation, we used the HT29 CRC cells that differentiate into the goblet cell lineage at confluence. We found that IκBα deletion (KO) by CRISP-Cas9 precluded goblet cell differentiation of human HT29 CRC cells at 7 days of post-confluence as determined by qPCR (Figure 5A) and WB analysis (Figure 5B) of the terminal differentiation markers MUC5AC and SPDEF.

**Figure 5.**
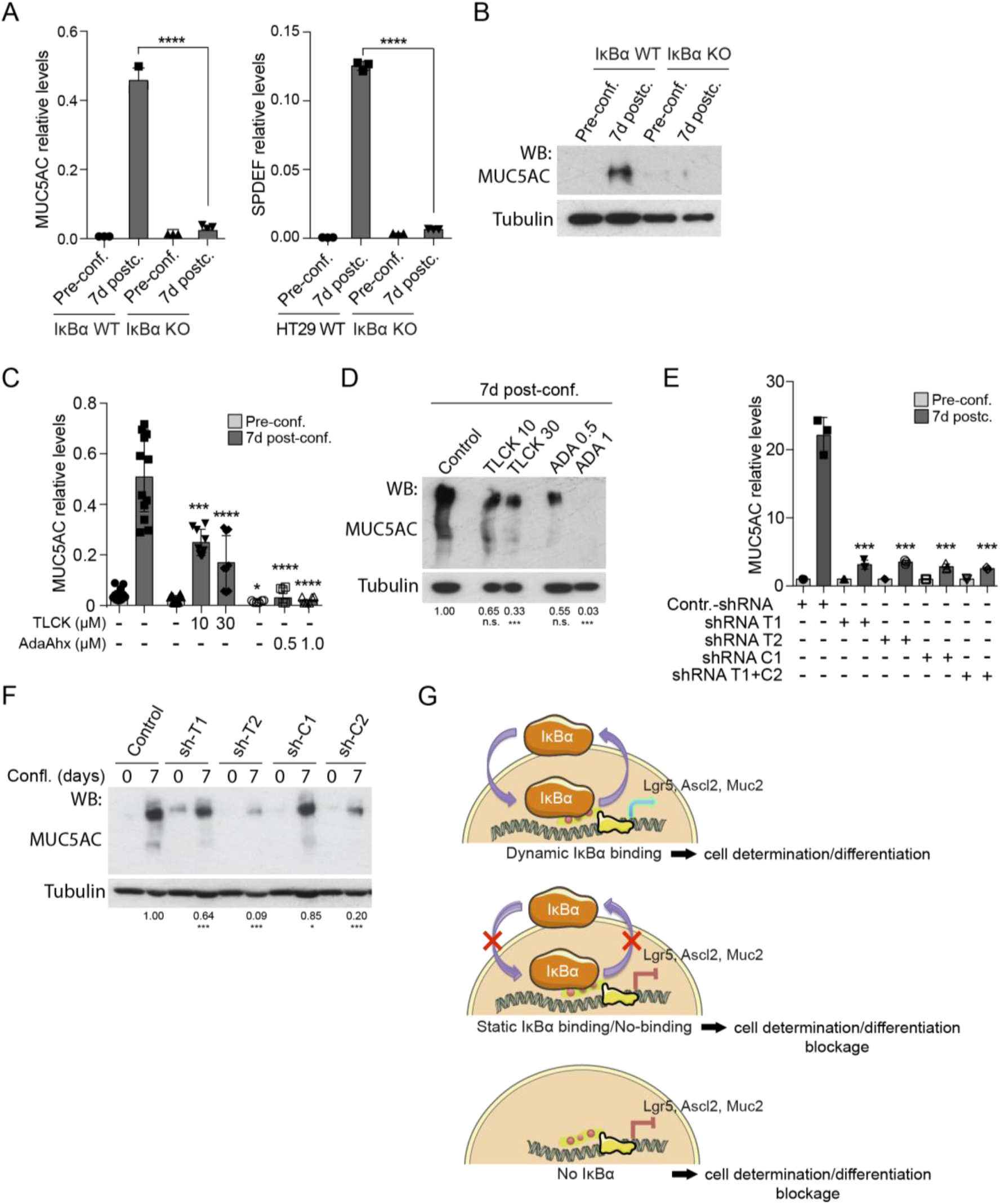
Functional impact of IκBα chromatin binding dynamics in cell differentiation. (A, B) qPCR (A) and WB analysis (B) of the indicated genes or proteins in parental or IκBα KO HT29 cells obtained at pre-confluence or 7 days post-confluence. (C, D) qPCR (C) and WB analysis (D) of MUC5AC in HT29 cells treated with the indicated protease inhibitors. (E, F) qPCR (E) and WB analysis (F) of MUC5AC in HT29 cells transduced with sh-RNAs against trypsin (T1, T2) or chymotrypsin (C1, C2). (G) Model for gene regulation by chromatin-associated IκBα. In brief, dynamic dissociation of IκBα from the chromatin at specific genetic loci promotes transcription of genes involved in stem cell maturation or differentiation (upper panel). Absence of dynamic binding (middle panel) or cells IκBα deficient (lower panels) fail to activate IκBα-dependent transcription thus imposing a differentiation/maturation blockage. Images in A-F are representative of three independent replicates performed. *p* values were derived from unpaired two-tailed *t*-test, ***p-value<0.001, **p-value<0.01, * p-value<0.05, n.s. no significant.

We then tested whether inhibition of histone cleavage, which favors pS-IκBα chromatin retention, affected HT29 differentiation. Treatment with the trypsin-like inhibitors TLCK or AdaAhx_3_L_3_VS (Figure 5C and 5D) or knocking down trypsin or chymotrypsin with different shRNA (Figure 5E, 5F and S3) both precluded goblet cell differentiation of HT29 cells, comparable to the effects imposed by IκBα deletion. These results suggest that histone cleavage, which favors dissociation of IκBα from the chromatin, is required for intestinal cell differentiation.

Together our results support a model in which dynamic binding/dissociation of IκBα to/from chromatin regulates specific gene transcription, which is essential for stem cell maturation and subsequent progression towards tissue-specific mature lineages (Brena et al., 2020; Marruecos et al., 2020; Mulero et al., 2013 and this work). Thus, in the absence of chromatin-bound IκBα or in case of irreversible IκBα chromatin binding (i.e. after trypsin or chymotrypsin inhibition), tissue stem cells and/or progenitors are retained into an immature state and fail to express differentiation markers such as epidermal *Filaggrin* and *Keratin 10* (Mulero et al., 2013) or intestinal *Muc2*, *Lyz1* (Marruecos et al., 2020) and *Muc5* (this work) (see model in Figure 5G).

## Discussion

We have previously demonstrated the existence of moonlighting functions for the essential NF-κB inhibitor IκBα as modulator of PRC2 activity at specific genomic loci. Now, we show that nuclear IκBα preferentially binds acetylated H4 species that are restricted to the stem cell compartment of the intestine and skin where nuclear p-IκBα is also localized. Loss of acetylated histone H4 in the differentiated compartments is likely due to the cleavage of its N-terminal tail (the region substrate of histone acetylation) by the action of specific proteases. We have identified trypsin and chymotrypsin as the most likely proteases involved in histone H4 cleavage in vivo. Moreover, different modifications at K residues affect protease activity on histone H4, as is the case of the H4K5,8,12,16A mutation or the hyperacetylated H4K5,8,12,16Ac, as expected. This result, together with the observation that crypt-derived histone H4 is cleaved after incubation with soluble villus lysates, indicate that histone H4 in the crypts is not hyperacetylated but different H4Ac species (i.e. H4K5Ac, H4K8Ac, H4K12Ac) may coexist in this cellular compartment.

N-terminal cleavage of histones, mainly H3, was previously reported in other systems associated with organism development and cell differentiation (Duncan et al., 2008; Kim et al., 2016; Melo et al., 2017; Vossaert et al., 2014). We propose that N-terminal loss at specific histones might represent a general but specific mechanism of gene regulation during development and tissue differentiation imposed by levels and activity of proteases recognizing specific histones and histone codes. IκBα dissociation from particular gene promoters (i.e. during cell differentiation) may be linked to local histone cleavage events that could be, in turn, controlled by acetylation or other PTMs. This dynamics of IκBα binding and dissociation would then trigger a switch in the repertoire of activators and repressors bound as specific gene promoters leading to a whole transcriptional response, which we will further investigate.

Because of the significant impact of nuclear IκBα in cell differentiation and stem cell maturation, and our previous data obtained in squamous cell carcinoma (Mulero et al., 2013), we speculate that IκBα chromatin binding and histone cleavage could similarly play a relevant contribution to tumorigenesis and tumor metastasis, which mainly depend on the stem cell-like capacity of tumor cells. Our prediction is that IκBα may represent a unique cellular tool for the integration of pro-inflammatory signals provided by the tumor stroma into a transcriptional program that impacts in cellular stemness. This IκBα function that facilitates the maintenance of stem cell homeostasis in response to inflammation or damage, would also execute pro-tumorigenic and pro-metastatic activities linked to inflammatory tumor microenvironments thus counteracting the positive effect of the immune system.

In brief, we have identified a previously unanticipated linked between histone cleavage and IκBα function during cell differentiation, which could have a relevant impact in the field of tissue regeneration, stemness and tumor metastasis.

## MATERIALS AND METHODS

### Animal Studies

WT mice and Lgr5^GFP-CreERT^ mice were from *The Jackson Laboratories.* In all procedures, animals were kept under pathogen-free conditions, and animal work was conducted according to the guidelines from the Animal Care Committee at the Generalitat de Catalunya. The Committee for Animal Experimentation at the Institute of Biomedical Research of Bellvitge (Barcelona) approved these studies.

Transgenic mice carrying Cherry-histone H4 were generated by transduction of a piggyback construct to ES cells.

### Cell lines and reagents

All cells were grown in Dulbecco’s modified Eagle’s medium (DMEM) [Invitrogen] supplemented with 10% fetal bovine serum (FBS) [Biological Industries]. Cells were grown in an incubator at 37°C and 5% CO2. Cells used in these studies were HEK-293T [ATCC Ref. CRL-3216], HT29 [ATCC Ref. HTB-38D] and HCT-116 [ATCC Ref. CCL-247]. Reagents used are the following: TLCK (Tosyl-L-lysyl-chloromethane hydrochloride) [Abcam ab144542] and AdaAhx3L3VS [Sigma-Aldrich 114802]. HT29 IκBα KO cells were generated by CRISPR-Cas9 using guides targeting exon 1.

### Villus/crypt-enriched fractionation

Intestine of WT mice was extracted. Villus was separated from crypts mechanically, as described previously (Marruecos et al., 2020). Then, crypts were purified by mechanical disaggregation and filtration in a 70 μm cell strainer.

### Cell transfection

We used Polyethylenimine (PEI) [Polysciences Inc. Ref. 23996] as a carrier vector following standard methods. In brief, we diluted 4μl PEI per μg of DNA in serum-free DMEM and incubated 5 min at room temperature. Then, we added the DNA and incubated the mix 20min at RT. Finally, we incorporated the PEI/DNA solution to the cell cultures.

### Pull down and peptide immunoprecipitation (IP) assays

PD assays were performed as previously described (Espinosa et al., 2003). Briefly, GST fusion proteins were incubated with lysates for 45 min in a rotary shaker at 4°C. When indicated, nuclear extracts were boiled at 98°C for 5 min in the presence of 1% SDS to disassemble pre-existing protein complexes and then neutralized in 1% Triton X-100. Precipitates were resolved in SDS-PAGE and analyzed by IB. For peptide IP, histone H4 peptides [Synpeptide CO LTD] were synthesized as biotinylated N-terminal and C-terminal amides. Peptides were incubated overnight at 4°C with the indicated cell extracts and precipitated with streptavidin-sepharose beads for 45 min.

### Cell fractionation and Western Blot (WB)

For soluble and chromatin separations, cells were lysed 1mM EDTA, 0.1mM Na-orthovanadate (Na3VO4), 0.5% Triton X-100, 20mM β- glycerol-phosphate, 0.2mM PMSF, protease inhibitor cocktail, in PBS for 20 min on ice and centrifuged at 13,000 rpm. Supernatants were recovered as the soluble fraction, and the pellets were lysed in Laemmli buffer (1x SDS-PAGE buffer plus β-mercaptoethanol (BME) [Sigma, Ref. M-3148]) or in 1%SDS PBS, sonicated and treated with 1% TritonX-100. For histone enriched fractions, cells were lysed in 10mM HEPES pH7.9, 1.5mM MgCl2, 10mM KCl, 0.5mM DTT, 1.5mM PMSF, 100mM sodium butyrate, protease inhibitor cocktail and 0.2mM H_2_SO_4_ for 30min on ice and centrifuged at 10080g. Then supernatants were dialyzed against PBS. Lysates were analyzed by Western blotting using standard SDS– polyacrylamide gel electrophoresis (SDS-PAGE) techniques. In brief, protein samples were boiled in Laemmli buffer, run in polyacrylamide gels, and transferred onto polyvinylidene-difluoride (PVDF) membranes [Millipore Ref. IPVH00010]. Membranes were incubated overnight at 4°C with the appropriate primary antibodies, extensively washed and then incubated with specific secondary horseradish peroxidase–linked antibodies from Dako [Ref. P0260 and P0448]. Peroxidase activity was visualized using the enhanced chemiluminescence reagent [Biological Industries Ref. 20-500-120] and autoradiography films [GE Healthcare Ref. 28906835]. Gels were stained with Coomassie (Brilliant Blue G-250 [Sigma Ref.6104-58-1].

### Antibodies used

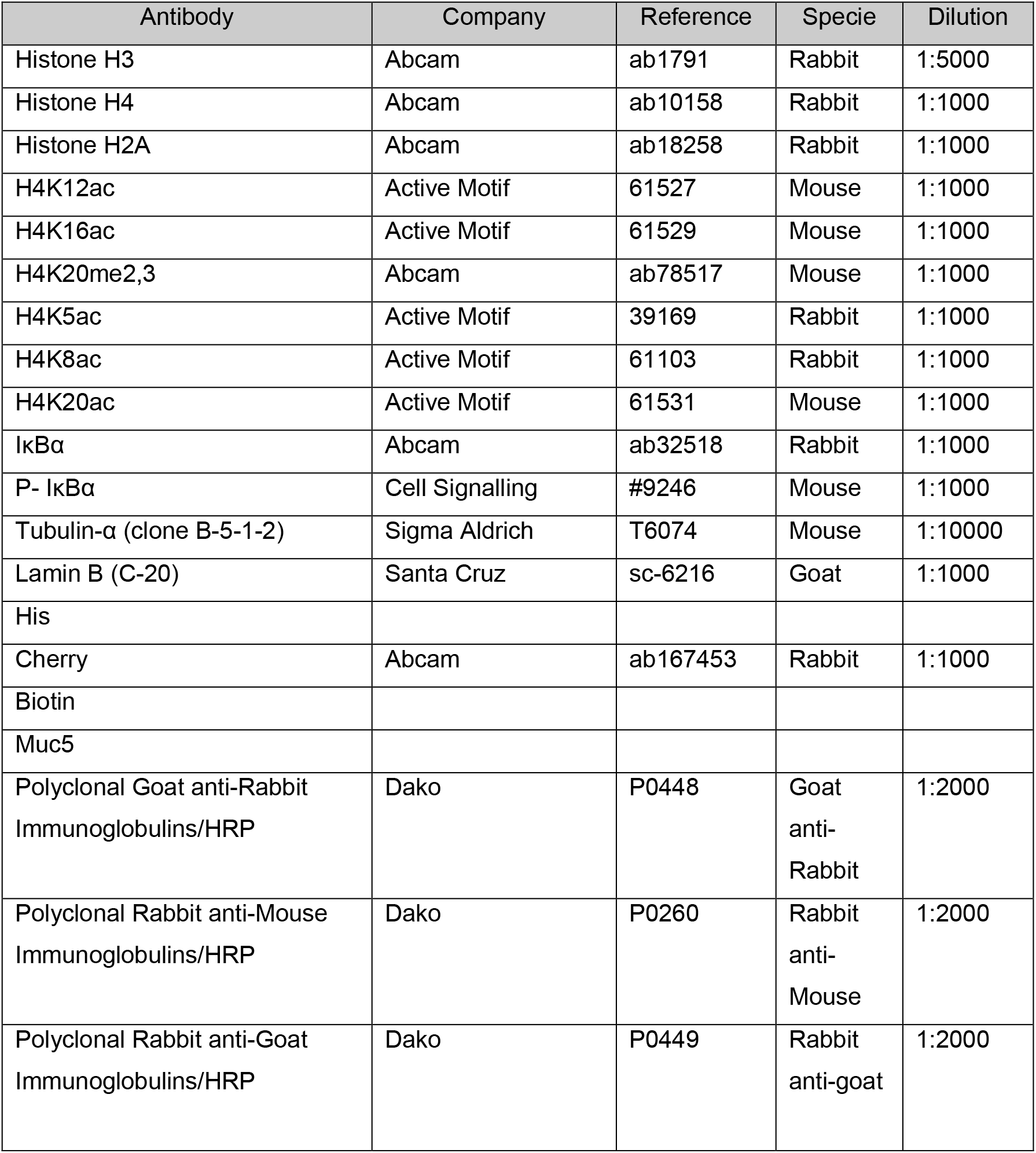

### Immunofluorescence (IF) analysis

Tissues were fixed in 4% formaldehyde overnight at room temperature and embedded in paraffin. 4μm paraffin embedded sections were first de-paraffinized in xylene. IHC was performed following standard techniques with EDTA- or citrate-based antigen retrieval and developed with the Envision+ System HRP Labelled Polymer anti-Rabbit [Dako Ref. K4003] or anti-Mouse [Dako Ref. K4001] and developed with TSA™ Plus Cyanine 3/ Fluorescein System [PerkinElmer Ref. NEL753001KT] and mounted in ProLong™ Diamond Antifade Mountant plus DAPI [Thermo Scientific Ref. P36971]. Images were taken in an SP5 upright confocal microscope (Leica).

### Size exclusion chromatography

GST-H2A or His-IκBα 1-200 proteins were expressed in BL21 bacteria and lysed as explained above. Then, proteins were purified using glutathione-Sepharose [Amersham Biosciences] or Nickel-NTA [Qiagen] resins, respectively. After elution, proteins were buffered exchanged using desalting columns [GE Healthcare] to a buffer containing 25 mM Tris pH 7.4, 150 mM NaCl, 0.2 mM EDTA pH 8.0, 1 mM PMSF and 1 mM DTT. Then, individual proteins (GST-H2A and His-IκBα 1-200) or a mix of both proteins at a ratio of 1:1 incubated at room temperature for 1hr to form complexes prior size exclusion chromatography were analyzed. 150 μl of each sample was loaded onto an analytical Superdex SD200 10/300 column [GE Healthcare] pre-equilibrated with the sample buffer and resolved at a flow rate of 0.5 ml/min on an AKTA Purifier [GE Healthcare] automated liquid chromatography system. 0.5 ml size fractions were collected and an equal volume aliquot of each fraction was analyzed by SDS-PAGE followed by Coomassie Blue staining.

### Recombinant histone octamer formation

pET11a-H2A, pET11a-H2B and pET11a-H3-H4 plasmids were a kind gift from Dr. James Kadonaga, UCSD. The original pET11a-H2B plasmid was mutated to pET11a-Ile-H2B (inserting an Isoleucine residue before the H2B sequence) to increase its protein yield. All histones were expressed in *Escherichia coli*BL21(DE3) by growing cells to A600 0.5-0.6 followed by induction with 0.2mM isopropyl β-D-1-thiogalactopyranoside (IPTG) for 3 h at 37°C.Cells were pelleted by centrifugation at 3000 rpm for 20 min at 4°C and washed once with phosphate-buffered saline (4mM Na_2_HPO_4_, 1mM KH_2_PO_4_, 137mM NaCl and 3mM KCl) (100 ml/liter bacterial cell culture). The H2A-H2B dimers and H3-H4 tetramers were purified following the detailed protocol previously published (Levenstein and Kadonaga, 2002).

Histone octamer formation was performed following the protocols from Dyer et al (Dyer et al., 2003) with some modifications. Briefly, histones molar ratio was quantified using A280 (Nanodrop) and applying an extinction coefficient for H2A/H2B,10.240 cm^−1^ M^−1^and for H3/H4, 9080 cm^−1^ M^−1^, respectively. Mixed histones were unfolded with Unfolding Buffer (20mM Tris-HCl pH 7.5, 10mM DTT and 7mM guanidinium hydrochloride) for 2h at 4°C. Then, the sample was dialyzed against the Refolding Buffer (10mM Tris-HCl pH 7.5, 1mM EDTA, 5mM BME and 2 M NaCl) at 4°C two times 3hr each and then ON. The day after, the sample was concentrated using a Centriprep 30-kDa cutoff membrane concentrator unit [Millipore] at 4°C and injected in a gel filtration analytical Superdex SD200 column. Fractions containing the histone octamer were pooled and dialyzed against Refolding Buffer with 50 % glycerol (10mM Tris-HCl pH 7.5, 1mM EDTA, 5mM BME, 2M NaCl and 50% (v/v) glycerol) at 4°C ON. Histone octamers were aliquoted and stored at −80C.

### Nucleosome formation

Original 601-DNA 12 copies plasmid was a kind gift from Dr. Karolyn Luger. Nucleosome core particles (NCP) were reconstituted following the classical salt gradient method. Briefly, we mixed 1ug of DNA with different volumes of histone octamers in a buffer containing 10mM Tris-HCl pH 8.0, 2M NaCl and 1mM EDTA. Samples were dialyzed using a 3.5 kDa cut-off Slide-A-Lyzer Mini dialysis device [Thermo Fischer Scientific]. We dialyzed the NaCl concentration from the initial 2M to 1.5M, 1M, 800mM, 600mM in the same buffer changing the dialysis every 3hr at 4°C. Finally, we left the dialysis going overnight at 2.5mM NaCl. The day after we changed the buffer one more time 3hr at 4°C in presence of 2.5mM NaCl. Successful NCP reconstitution was confirmed by running a native electrophoretic mobility shift assay (EMSA) loading 10 ng of NCPs in a 4% native acrylamide gel (19:1) in the cold room. Before loading, samples were mixed with 10% (v/v) glycerol in absence of bromophenol blue to preserve the integrity of the NCPs.

### Non-radiolabeled electrophoretic mobility shift assay

Purified full length His-IκBα protein was incubated with PKA in the kinase buffer 20mM HEPES pH 7.9, 150mM NaCl, 10mM MgCl2, 10mM NaF, 0.2mM sodium orthovanadate and 20μM ATP for 30 minutes at 30°C. Immediately after, NCP were incubated with increasing concentrations of phosphorylated IκBα (p-IκBα) for 1hr at 4°C. p-IκBα was obtained by incubation of recombinant IκBα with active IKKß in the presence of ATP. When indicated, anti-IκBα antibody was added to the mix to block the NCP-IκBα interaction. After, NCP-IκBα complexes were run in a 4% native acrylamide gel (19:1) in the cold room. Gels were stained using GelRed [Biotium] for 30 min and destained using TBE 0.5× 3 times for 10min in the cold room. Gels were imaged using a GelDoc system [Bio-Rad].

### Radiolabeled electrophoretic mobility assay

601-DNA was radiolabeled with ^32^P using T4-polynucleotide kinase and [γ-^32^P] ATP for 1hr at 37°C and then incubated with the histone octamer in presence of 2M NaCl. Here, NCP formation was achieved by sequential dilution instead of dialysis as described above. Radiolabelled NCPs were incubated with the proteins under study for 20 min at room temperature in binding buffer 10mM Tris-HCl (pH 7.5), 50mM NaCl, 10% (v/v) glycerol, 1% (v/v) NP-40, 1mM EDTA, and 0.1 mg/mL PolydIdC. Samples were run in TGE buffer (24.8mM Tris base, 190mM glycine, and 1mM EDTA) at 200V for 1hr, and the gel was dried. Protein complexes were analyzed by native electrophoresis on a 4% (w/v) native acrylamide gel.

### Knock down assays

MISSION shRNA for trypsin and chymotrypsin plus lentiviral plasmids were transfected in HEK293T cells. One day after transfection, media was refreshed. Virus was collected 24 hours later and then concentrated using Ultracentrifuge Optima™ XPN-100 - IVD (Biosafe) [Beckman Coulter]. Cells were infected and selected with puromycin.

### Chromatin Immunoprecipitation (ChIP) and ChIP-sequencing analysis

Human colon cell lines were subjected to chromatin immunoprecipitation (ChIP) as previously described (Aguilera et al., 2004; Mulero et al., 2013). Briefly, formaldehyde crosslinked cell extracts were sonicated, and chromatin fractions were incubated for 16 hours with anti-IκBα [abcam ab32518], anti- P-IκBα [Cell Signaling #9246], anti-Acetylated H4 [Abcam ab177790] and anti-H4K12ac [Active Motif 61527] antibodies in RIPA buffer and then precipitated with protein A/G-sepharose [GE Healthcare, Refs. 17-0618-01 and 17-0780-01]. Crosslinkage was reversed and 6-10 ng of precipitated chromatin was directly sequenced in the genomics facility of Parc de Recerca Biomèdica de Barcelona (PRBB) using Illumina HiSeq platform.

Raw single-end 50-bp sequences were filtered by their quality (Q>30) and length (length > 20-bp) with the Trim Galore software (available at http://www.bioinformatics.babraham.ac.uk/projects/download.html#trim_galore). Filtered sequences were aligned against the reference genome (mm10 release) with Bowtie2. MACS2 software was run first for each replicate, and then by combining all replicates, using unique alignments (q-value < 0.1). Broad peaks calling was set. Peak annotation was performed with ChIPseeker package and functional enrichment analysis with enrichR using the latest version of GO annotations. ChIP-sequencing data are submitted to GEO database.

### Cell sorting

Villus and crypt cells were obtained after mechanical disaggregation and 40μm filtration. Cells were incubated with APC-EphB2 antibody [BD Pharmingen Ref. 564699] for 20min, stained with DAPI and the sorted in an Influx™ Sorter [BD Biosciences].

### qRT-PCR analysis

Total RNA was extracted with the RNeasy Mini Kit [Qiagen Ref. 74004] and the RT-First Strand cDNA Synthesis Kit [GE Healthcare Life Sciences Ref. 27-9261-01], was used to produce cDNA. qRT–PCR was performed in LightCycler480 system using SYBR Green I Master Kit [Roche Ref. 04887352001]. The primers used are listed in the table below.

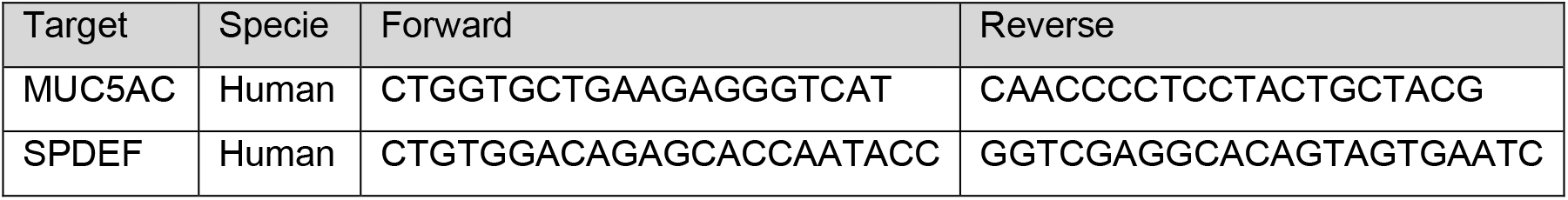

### RNA-sequencing experiments and data analysis

We extracted total RNA using RNeasy Micro Kit [Qiagen Ref. 74004]. The RNA concentration and integrity were determined using Agilent Bioanalyser [Agilent Technologies]. Libraries were prepared at the Genomics unit of PRBB (Barcelona, Spain) using standard protocols and cDNA was sequenced using Illumina® HiSeq platform, obtaining ~25 to 30 million 50bp single-end reads per sample. Adapter sequences were trimmed with Trim Galore. Sequences were filtered by quality (Q>30) and length (>20 bp). Filtered reads were mapped against the latest release of the mouse reference genome (mm10) using default parameters of TopHat (v.2.1.1) (Kim et al., 2013) and expressed transcripts were then assembled. High quality alignments were fed to HTSeq (v.0.9.1) (Anders et al., 2015) to estimate the normalized counts of each expressed gene. Differentially expressed genes between different conditions were explored using DESeq2 R package (v.1.20.0) (Love et al., 2014). Plots were done in R. RNA-sequencing data are deposited at the GEO database with accession number GSE131187.

### Quantification and Statistical Analysis

Statistical parameters, including number of events quantified, standard deviation, and statistical significance are reported in the figures and in the figure legends. Statistical analysis has been performed using GraphPad Prism6 software (GraphPad) and p<0.05 is considered significant. Two-sided Student’s t-test was used to compare differences between two groups. Each experiment shown in the manuscript has been repeated at least twice.

## Data availability

RNA-Seq data: GEO databaseGSE131187 https://www.ncbi.nlm.nih.gov/geo/query/acc.cgi?acc=GSE131187

ChIP-seq data has been submitted to GEO database.

## ACKNOWLEDGEMENTS

We want to thank the Bigas’ and Espinosa’s lab members for constructive discussions and suggestions and technical support. This work has been funded by Instituto de Salud Carlos III FEDER (PI19/0013), Agencia Estatal de Investigación, Spain (PID2019-104695RB-I00) to A.B., NIH/GM085490 to G.G.,

BIO2017-83650-P to JVF and Generalitat de Catalunya 2017SGR135. LM is a predoctoral fellow of 2015FI-B00806 and 2016FI-B1 00110 and MF has financial support by the Universitat de Vic-Universitat Central de Catalunya PhD fellowships program. The authors thankfully acknowledge the computer resources at Pirineus and the technical support provided by the Spanish Supercomputer Network (BCV-2020-1-0001).

## Supplemental Material

### SUPPLEMENTARY FIGURES AND LEGENDS

**Figure S1.**
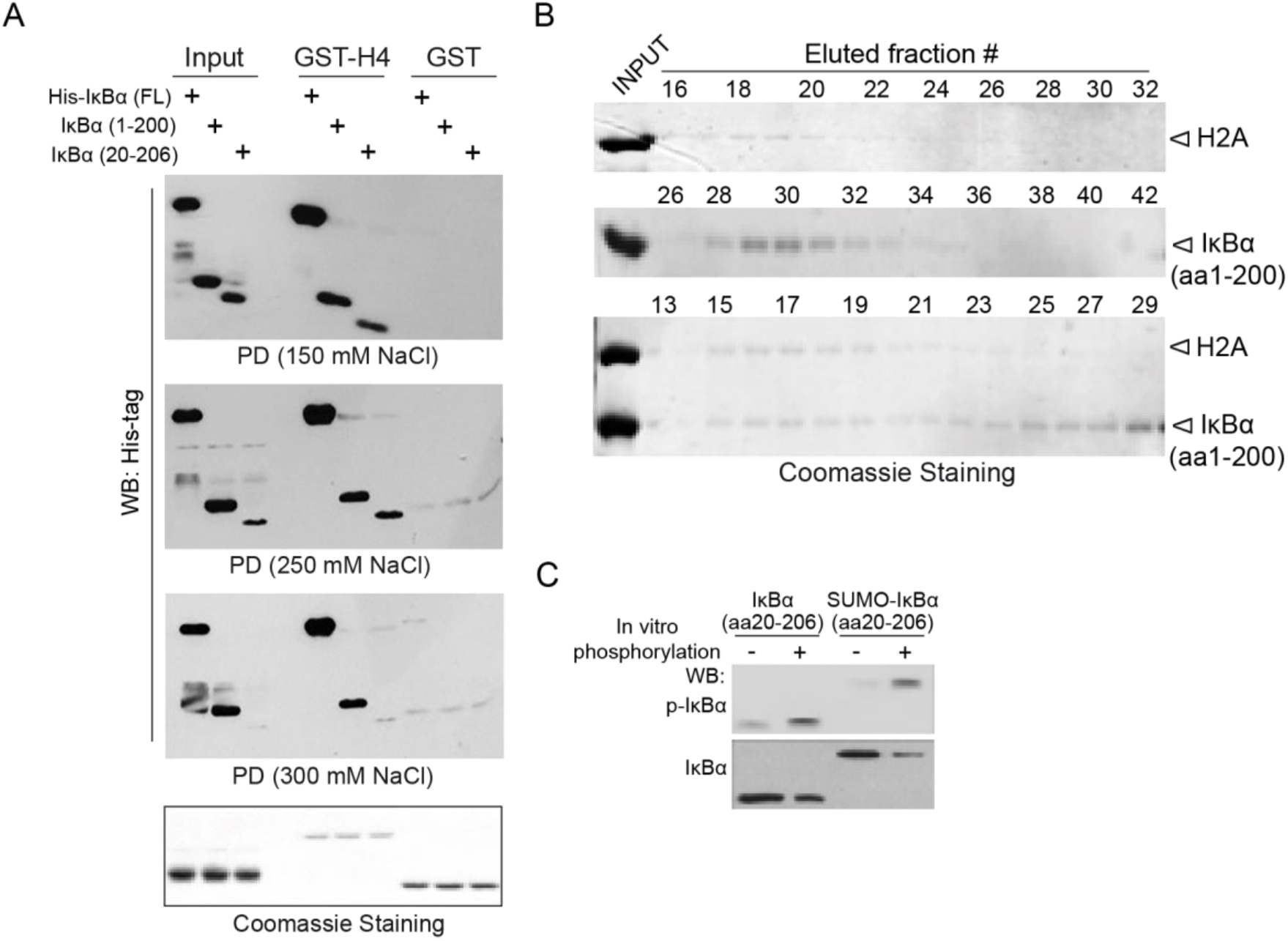
In vitro interactions and phosphorylation of IκBα. (A) PD experiments under different salt concentrations using GST-H4 as bait and the indicated IκBα constructs expressed in HEK-293T cells. (B) Coomassie staining analysis of the indicated fractions recovered in the <u>F</u>ast Protein <u>Li</u>quid <u>C</u>hromatography (FPLC) analysis of IκBα and histone H2A complexes. (C) Western blot analysis of IκBα phosphorylated in vitro by addition of active IKKβ kinase with anti-p-IκBα (S32-36) antibody.

**Figure S2.**
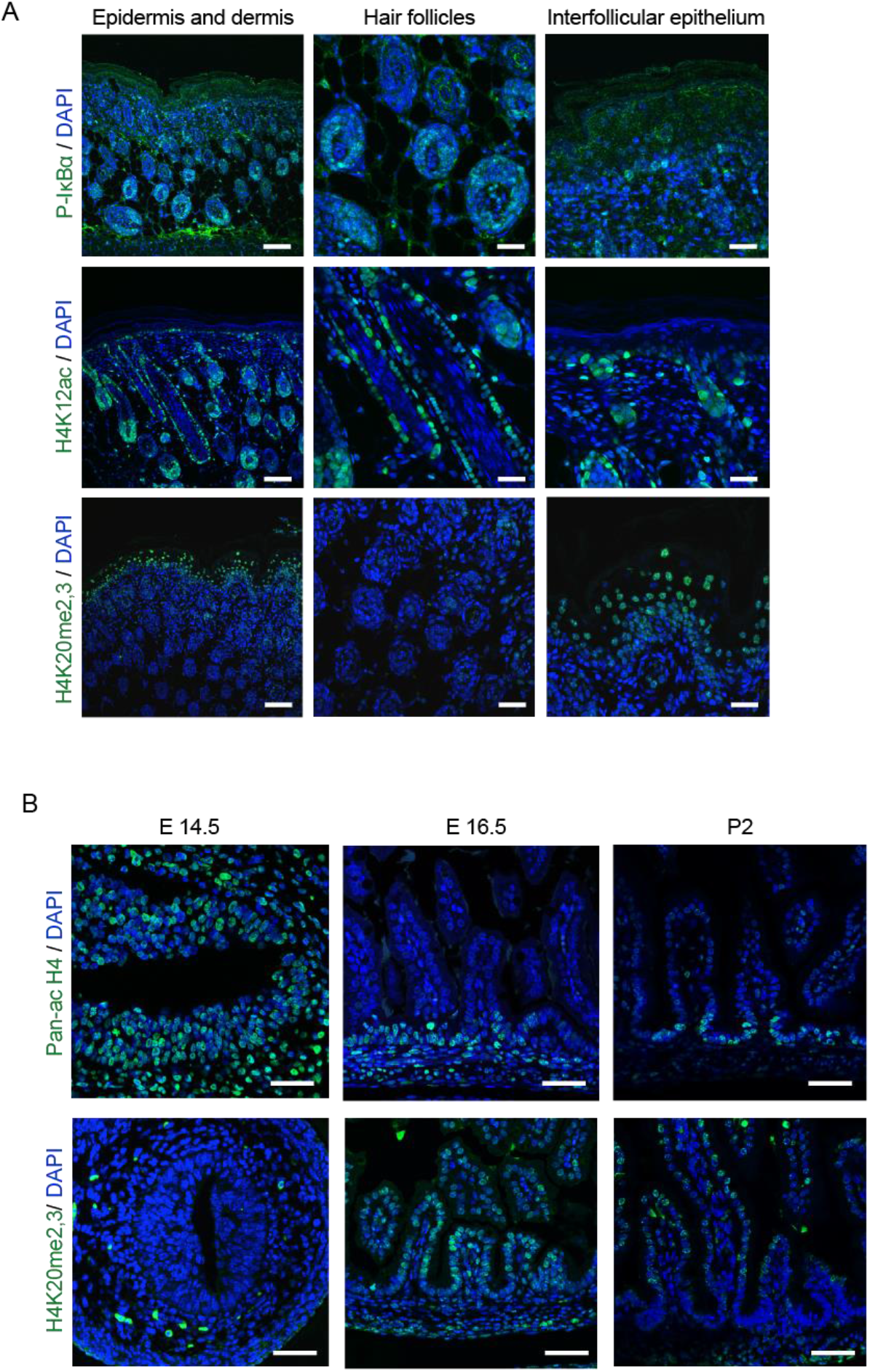
IκBα is localized in stem cell compartments. (A, B) IF analysis with the indicated antibodies in sections from murine skin of 2-month-old mice (A) and sections from small intestine of mice at different stages of development (B). Scale bars in A and B, 25μm.

**Figure S3.**
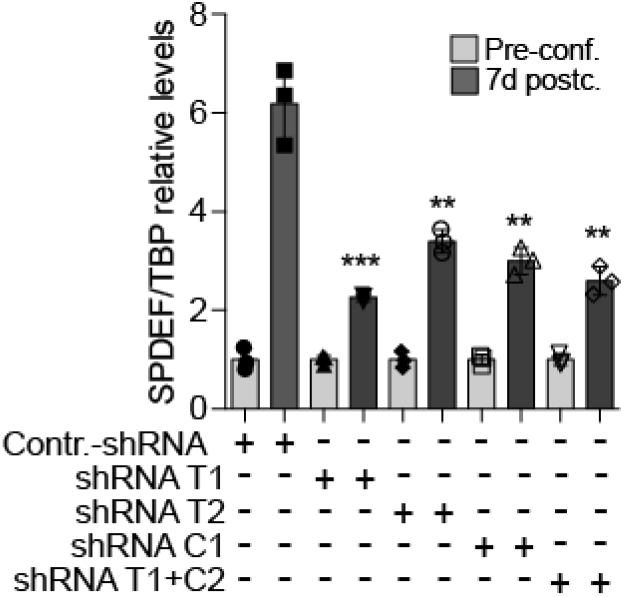
Inhibition of goblet cell differentiation by trypsin or chymotrypsin knock-down. Analysis by qPCR of SPDEF in HT29 cells transduced with specific shRNA against trypsin (T) or chymotrypsin (C) at pre-confluence or 7 days after confluence.

